# Mechanical properties of plasma membrane vesicles correlate with lipid order and viscosity and depend on cell density

**DOI:** 10.1101/669085

**Authors:** Jan Steinkühler, Erdinc Sezgin, Iztok Urbančič, Christian Eggeling, Rumiana Dimova

**Affiliations:** Theory and Bio-Systems, Max Planck Institute of Colloids and Interfaces, Science Park Golm, 14424 Potsdam, Germany; MRC Human Immunology Unit, Weatherall Institute of Molecular Medicine, University of Oxford, Headley Way, Oxford OX3 9DS, UK; Condensed Matter Physics Department, “Jožef Stefan” Institute, Ljubljana, Slovenia; Institute of Applied Optics Friedrich-Schiller-University Jena, Max-Wien Platz 4, 07743 Jena, Germany; Leibniz Institute of Photonic Technology e.V., Albert-Einstein-Straße 9, 07745 Jena, Germany

## Abstract

Plasma membranes dynamically respond to external cues and changing environment. Quantitative measurements of these adaptations can elucidate the mechanism that cells exploit to survive, adapt and function. However, cell-based assays are affected by active processes while measurements on synthetic models suffer from compositional limitations. Here, as a model system we employ giant plasma membrane vesicles (GPMVs), which largely preserve the plasma membrane lipidome and proteome. From analysis of fluorescence emission and lifetime of environment-sensitive dyes, and membrane shape fluctuations, we investigate how plasma membrane order, viscosity and bending rigidity are affected by different stimuli such as cell seeding density in three different cell models. Our studies reveal that bending rigidity of plasma membranes vary with lipid order and microviscosity in a highly correlated fashion. Thus, readouts from polarity- and viscosity-sensitive probes represent a promising indicator of membrane mechanical properties. Quantitative analysis of the data allows for comparison to synthetic lipid membranes as plasma membrane mimetics.

## Introduction

Plasma membrane (PM) remodelling governs essential cellular processes such as endo- and exocytosis, cellular division, nanoparticle and viral uptake and vesicle shedding (1–3) and the energetic cost of elastic membrane deformation makes up a significant contribution to these processes (3). However, progress in the field is limited by lack of methods to examine the PM mechanical properties (4). Cellular complexity (in particular, the intertwinement of the plasma membrane with actin cytoskeleton and glycocalyx as well as its involvement in active processes such as endo- and exocytosis) poses a great challenge to measuring these properties in intact cells. To this end, model membrane systems such as giant unilamellar vesicles are applied. However these synthetic systems lack the compositional complexity of the cellular plasma membrane. Here, we show that isolation and study of PM blebs bridge the gap between mechanical manipulation of living cells and assays based on the use of synthetic cell-sized membranes, as exemplified by giant unilamellar vesicles (5–7). After the initial reports (8) about isolated PM blebs or giant plasma membrane vesicles (GPMVs), they have gained significant attention in the field of liquid-liquid phase separation in the membrane (9, 10). However, up to now they have rarely been used for the study of the PM mechanical properties (11). An interesting aspect of GPMVs as a PM model is their compositional and structural complexity, closely representing the PM lipidome and proteome (12, 13), and the preserved lipid-protein (14) and protein-protein (15) interactions. As GPMVs are completely free of cytoskeletal support (16), they can be used to distinguish the contribution of the PM from that of active processes or the cytoskeleton in mechanical manipulation assays (such as micropipette aspiration and tether-pulling) on living cells (17–19). In this way, GPMVs are ideally suited to address emerging questions of the role of lipids and lipid-protein assemblies in mechanical and structural cellular response. Here, we use GPMVs to study the PM mechanics, membrane packing and viscosity while subjecting the cells to a range of different stimuli. By applying a spectrum of established characterization methods to GPMVs, we compare the readout from the used probes in PM and synthetic membranes and asses the correlation of molecular order and viscosity with membrane mechanics.

## Results and Discussion

### Bending rigidity, lipid packing and viscosity measurements

The bending energy of lipid bilayers are well described by the elastic sheet model with a single parameter, namely the bending rigidity *κ*, which takes values on the order of 20-30 k_B_T for single-component membranes made of typical phosphocholines (20). Bending rigidities reported from tube-pulling measurements on cells appear to be a factor of 2-10 higher (19), but the detected presence of actin and other cytoskeletal material in the pulled tubes is bound to influence the results. The bending rigidity of blebs bulging out from adhering cells was found strongly dependent on whether the bleb was expanding (softer and closer in bending rigidity to model lipid membranes) or retracting (stiffer) (21). In the GPMVs studied here, the membrane (and the whole vesicle) is detached from the cytoskeleton. The GPMVs were isolated from adherent U2OS cells by chemically induced cleavage of the cytoskeleton from the PM (22) (see Methods for details). When observed at room temperature using optical microscopy, U2OS GPMVs are mostly spherical and exhibit high optical phase contrast. Staining them with membrane dyes leads to homogenous dye distribution with no observable phase separation at room temperature (**Fig. 1 a,b**). After slight osmotic deflation, membrane fluctuations are readily observed. Membrane fluctuations around the mean spherical shape were analysed (following the approach in (23); see **Fig. 1b** red line) and a power spectrum of the amplitude and energy of each fluctuation mode was obtained (**Fig. 1c**). The power spectrum deduced from the elastic sheet model (see Eq. 1 in Material and Methods) was fitted with good agreement to the experimental data within the limiting factors of optical resolution and noise which become apparent at higher mode numbers (high q values). Thus, we conclude that GPMV bending deformations up to length scales of the optical resolution (0.5 μm) can be described with a single parameter - the bending rigidity. It is worth noting that the bending rigidity is a material property of the membrane, while the membrane tension (which can be also assessed from fluctuation analysis) depends on the vesicle state and the forces acting on the membrane. In contrast to the more complex conditions in cells (24, 25), the membrane tension of GPMVs should be simply set by the area-to-volume ratio, or more broadly speaking, the shape of the vesicle. GPMVs used for fluctuation analysis were osmotically deflated and hence exhibited a low membrane tension of about 0.01 μN/m (which is similar in order of magnitude to that measured in cells (17)). As can be expected, the tension was not observed to vary with GPMV isolation conditions (Fig. S1).

**Figure 1.**
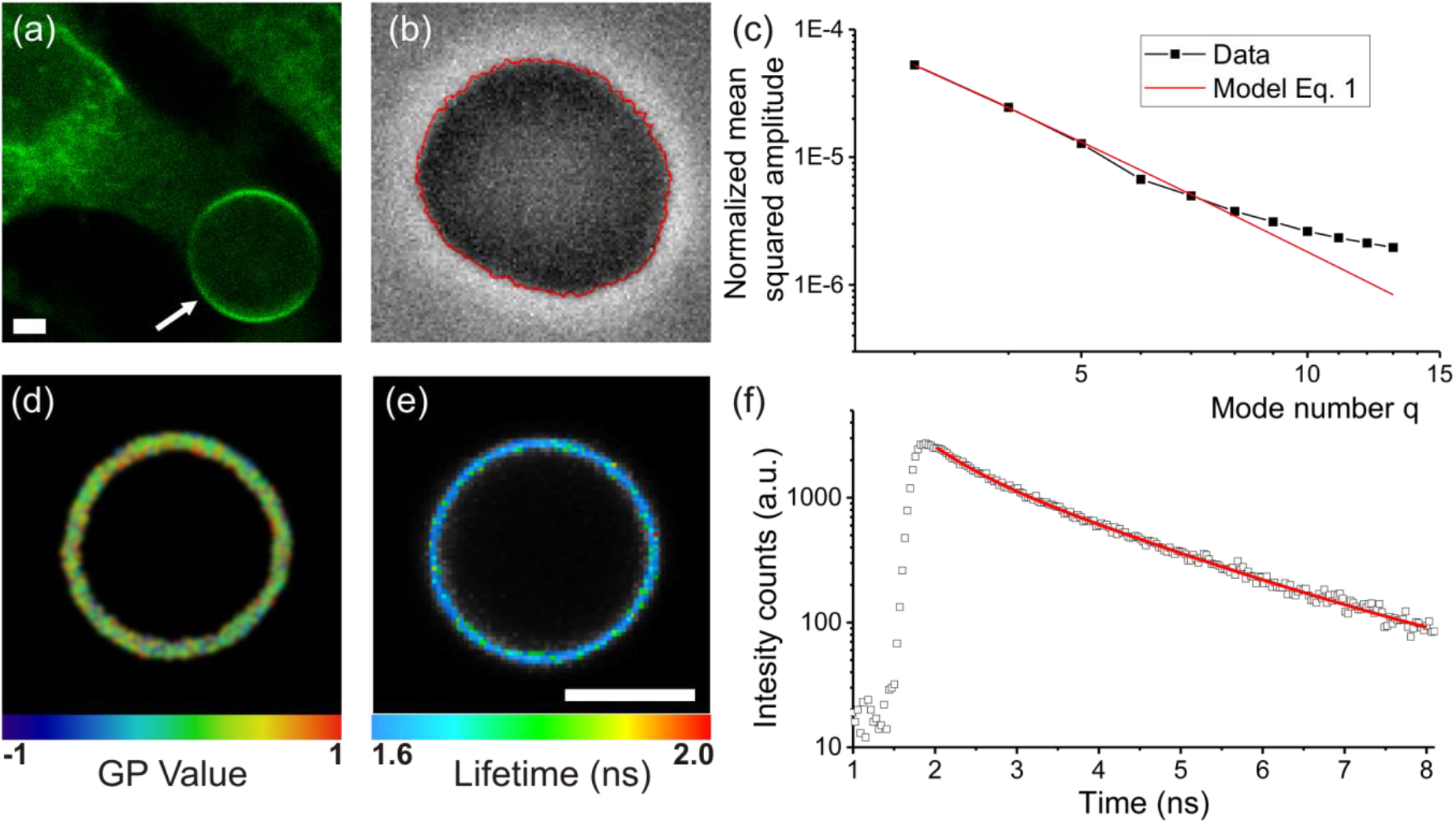
**(a)** GPMV formation. Chemically induced cytoskeletal cleavage of the plasma membrane results in the formation of vesicles (arrow), which subsequently detach from the cell (confocal cross-section). All experiments were conducted at room temperature where GPMVs exhibit one liquid phase as seen by homogenous Fast-DiIC18 (green) distribution, the variation of brightness along the contour is due to optical polarization effects. **(b, c)** Bending rigidity measurements by analysis of thermally-induced membrane undulations. Detected membrane contour (red) is shown overlaid on the phase contrast image in (b). From the contour analysis, the power spectrum (black data in (c)) is obtained and fitted to Eq. 1 (red curve). **(d)** GP measurement on a GPMV membrane. Color code corresponds to extracted GP map. **(e, f)** Fluorescent lifetime of molecular rotor embedded in the GPMV membrane. Color code in (e) indicates average fitted rotor lifetime. Lifetime histogram from the whole membrane region of a GPMV (open points) and the biexponential fit to the data (red curve) are shown in (f). All scale bars indicate 5 μm.

To correlate the continuum mechanical properties with molecular-scale descriptors, PM order and viscosity were also examined. Measurements were conducted at room temperature, above the temperature of phase separation of GPMVs (26), to ensure homogeneous membrane properties across each vesicle. The lipid order was assessed from the spectral response of a polarity-sensitive membrane probe C-Laurdan, which exhibits a red-shifted emission in more disordered membranes (27, 28). For easier comparison, local spectral properties were converted to the so-called generalized polarization parameter (GP, **Fig. 1d**), which represents a relative index for molecular lipid packing. GP ranges between +1 and −1, with higher values representing higher lipid order (27, 29). To further interrogate the local PM viscosity, GPMVs were stained with a Bodipy-based molecular rotor (30, 31). The probe lifetime as explored with fluorescence lifetime imaging microscopy (FLIM) is sensitive to the membrane viscosity. **Fig. 1e** shows an exemplary FLIM map, generated pixel-wise from respective histograms (**Fig. 1f**) of photon arrival times. This combination of methods was used in the next sections to obtain a detailed characterization of GPMVs isolated in varying conditions.

### Influence of GPMV isolation chemicals

GPMVs can be prepared using various conditions (8), but it is unclear how the different approaches affect the membrane mechanical properties. Modulation of elastic properties might be anticipated because changes in lipid phase by varying extraction methods are well documented. For example, GPMVs prepared with dithiothreitol (DTT) and paraformaldehyde (PFA) exhibit a demixing temperature about 10-20°C higher than that of GPMVs derived using N-Ethylmaleimide (NEM) (26). DTT has been shown to affect lipid-lipid and lipid-protein interactions and to integrate directly into lipid membranes (32, 33). To figure out potential differences in mechanical properties, the bending rigidity of GPMVs formed using these two most widely employed methods was assessed. GPMVs prepared by incubation with DTT+ PFA or NEM show only a small (<2*k*_*B*_*T*) difference in bending rigidity (**Fig. 2a**). Apparently, the artefacts induced by DTT do not exhibit a strong effect on the bending rigidity.

We also questioned the effect of PFA (34), which may be potentially inducing (membrane) protein crosslinking, even when used at much lower concertation then in cell fixation protocols. GPMVs were extracted using NEM, and 25 mM PFA was added to the GPMV suspension followed by 1 hour incubation. No significant influence of PFA on the bending rigidity was observed, indicating that potential effects of protein-protein crosslinking are small when considering the bending rigidity variance between individual GPMVs (**Fig. 2a**).

**Figure 2.**
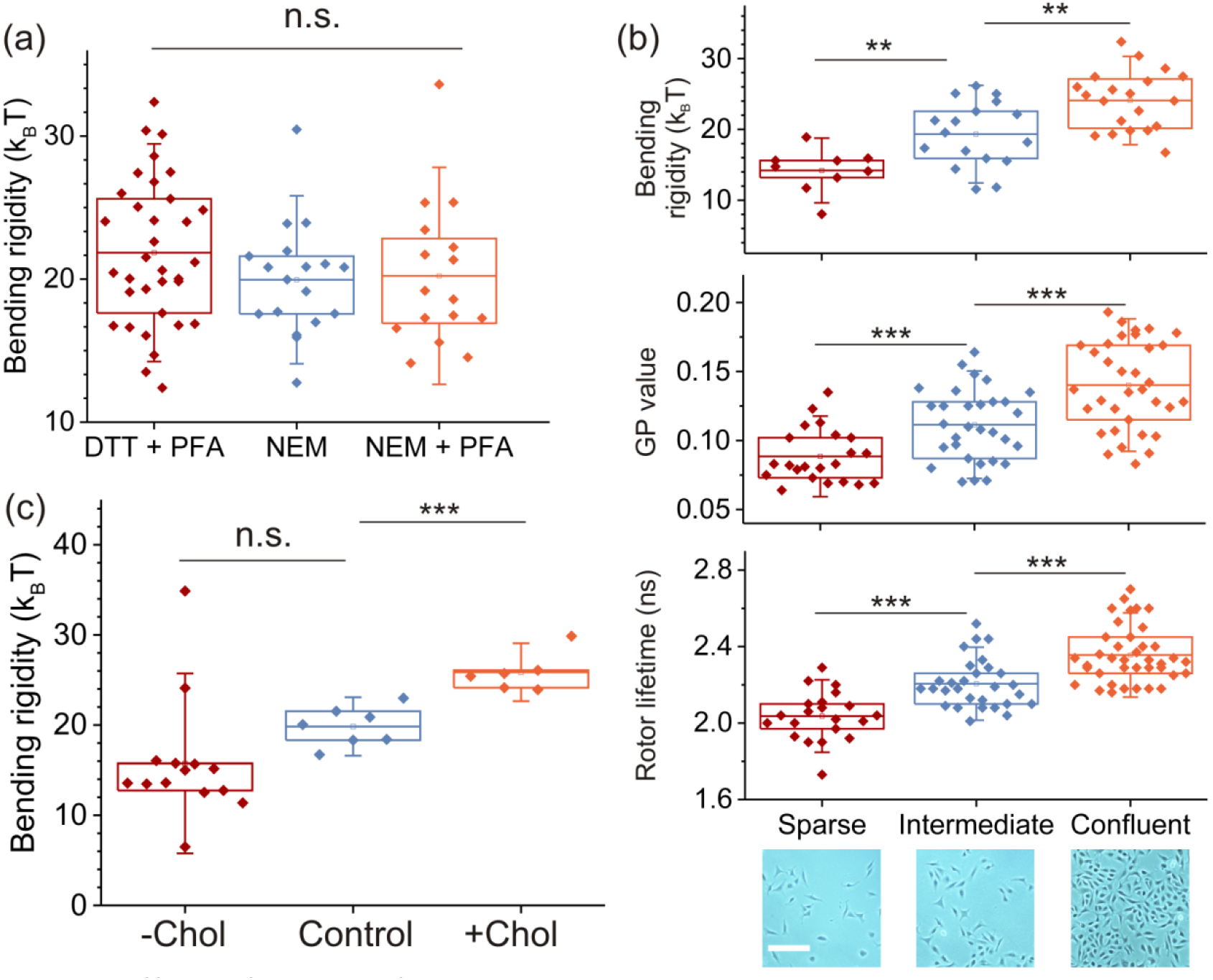
Effect of cell confluency, isolation chemicals and cholesterol content on the mechanical and molecular properties of PM membranes. **(a)** GPMV bending rigidity values for isolation using 2 mM DTT+ 25 mM PFA, 2 mM NEM, and 2 mM NEM with subsequent addition of 25 mM PFA. **(b)** GPMV bending rigidity, GP and rotor lifetime for varying cell densities at time of GPMV isolation by incubation with 2mM DTT + 25mM PFA; (bottom) phase contrast images (scale bar: 200 μm) of adherent cells before isolation shown below. **(c)** Effect of cholesterol extraction (-Chol), enrichment (+Chol) or buffer only (control) treatment of cells before GPMV extraction. Double-sided Student p-test was used to estimate significance with n.s. p> 0.05 **p<0.01, *** p<0.001. Each data point indicates one individual vesicle. Boxes have the conventional meaning of lower 25% and 75% quartile around the population mean value (middle line) and error bars indicate 1.5 std. dev.

### Cell density modulates membrane properties

Adherent cells have been reported to adapt their membrane composition depending on the local cell density and time in culture after seeding (35–37), but it remains unclear how PM elastic response is related to changes in membrane composition. We extended our studies to GPMVs isolated from cells at varying confluency, but constant time in culture. We found a strong correlation between initial cell density and GPMV bending rigidity, with a trend for a stiffer phenotype at higher cell confluency (**Fig. 2b top**). Interestingly, PM stiffening has been related to reduced cell migration (11). The bending rigidity measured here for cells in sparse conditions are consistent with those reported for blebs expanding from adhering cells (21). Note that significantly higher values for the bending rigidity of cell membranes have also been reported (19), but these data were obtained from tube-pulling experiments where the detected presence of actin in the pulled tubes could explain the difference.

To further understand the relation between GPMV membrane physical properties and confluency, we assessed the PM order and viscosity, which were found to increase with cell confluency (**Fig. 2b middle and bottom**). These results are consistent with the finding that growth conditions modulate the phase transition temperature of GPMVs derived from cells at varied confluency (38). We find that in the GPMV model system, molecular membrane properties such as lipid order (reported by the C-Laurdan GP value) and dynamic parameter of microviscosity (assessed from lifetime of Bodipy rotor) correlate with the equilibrium elastic parameter of bending rigidity (**SI** **Fig. S2**).

### Cholesterol content and temperature modulate PM bending rigidity

It is already well established that PM order (assessed by GP value of polarity sensitive membrane probes, such as C-Laurdan) is related to cholesterol content in cell PMs and GPMVs (39, 40). Thus, we questioned whether the bending rigidity of derived GPMVs is related in similar fashion to cellular PM cholesterol content. To this end, cholesterol concentration in the PM was directly altered by incubation of the cells with methyl-β-cyclodextrin before GPMV isolation; cyclodextrins are agents actively used for cholesterol exchange (41, 42). In the used conditions, about 60% cellular cholesterol was extracted from the PM (43). GPMVs isolated from cholesterol-depleted cells (−Chol) appear softer while cholesterol enrichment (+Chol) stiffens GPMV membranes (**Fig. 2c**). It is tempting to speculate that the bending rigidities of cholesterol depleted and enriched GPMVs exemplify the respective bending rigidities of liquid ordered Lo (“lipid-raft like”) and liquid disordered Ld phases in the PM. For their ratio we find κ_Lo_/κ_Ld_≈1.7. This stiffness mismatch is consistent with the membrane morphologies we observe in GPMVs exhibiting coexisting Lo-Ld phases (see **Fig. S3**): to minimize the bending energy, the lipid raft-like phase exhibits lower membrane curvature and curvature stresses preferably act on the softer non-raft phase bending it in the vicinity of the two-phase contact line, similarly to behaviour observed in L_o_-L_d_ phase separated synthetic model membranes (44, 45).

Apart from cholesterol content, membrane fluidity also changes with temperature. Generally, all experiments were conducted at room temperature (23°C). However, when bending rigidity experiments were performed at physiological temperature, GPMVs were found to soften from κ(23°C) ≈ (5.1 ± 1.2) 10^−20^ J (data from **Fig. 3a**) to κ(37°C) ≈ (2.3 ± 1.2) 10^-−20^ J (n=6). This result is again consistent with reduction of membrane order (C-Laurdan GP value) at increased temperature (34, 46). Given these lines of evidence, we aimed to further explore the interdependence of lipid order (C-Laurdan GP value) and bending rigidity.

**Figure 3.**
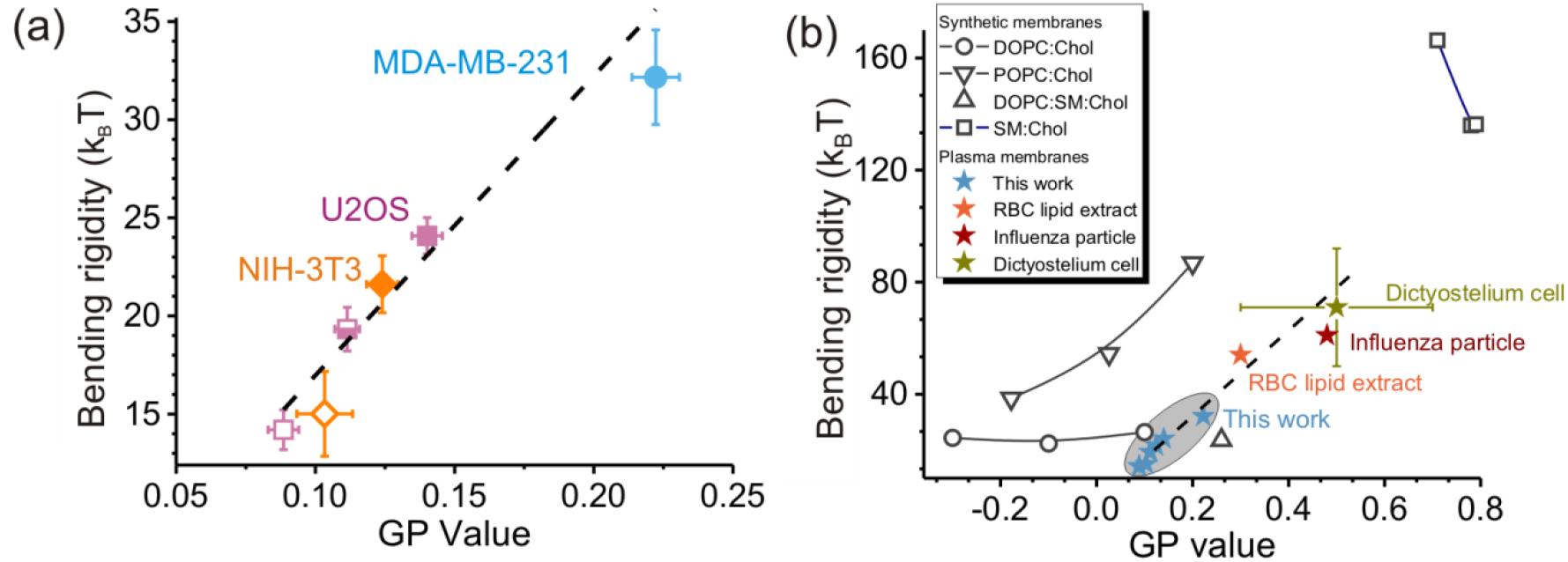
Correlation between plasma membrane lipid order, reported by C-Laurdan GP, and bending rigidity and comparison to lipid-only membranes. **(a)** Correlation between mean values of C-Laurdan GP, bending rigidity obtained on GPMVs harvested from different cell lines (as indicated by the colour) and at varying cell density (as indicated by the filling of the symbol: open – sparse, half-filled – intermediate, solid – confluent). The relation between GP values and the bending rigidity κ is well approximated by κ(GP) = (153*GP + 2)*k*_*B*_*T* (adj. R^2^ ≈ 0.91) shown as dashed line. **(b)** Literature values for C-Laurdan GP and bending rigidity of DOPC:Chol (open circles) with increasing fraction of cholesterol from left to right 1:0, 7:3, 5:5 molar ratios, POPC:Chol (triangles) 1:0, 9:1, 2:1, DOPC:SM:Chol (solid square) 7:1:2, SM:Chol (open blue squares) 9:1, 8:2, 7:3. Data from lipid vesicles (black symbols), RBC lipid extract extract (orange star), influenza particle (cherry) and Dictyostelium (ochre) are adapted from (20, 23, 47–54). Dictyostelium PM bending rigidity was estimated from talin-deficient cells (55). Blue stars indicate values found for GPMVs at varying confluency in this study (same data as in panel a). Errors bars (std. error) are only shown if known and are significantly larger than the size of the data point (panel b). The dashed line is identical to that in panel a.

### Membrane order and bending rigidity correlate in a wide range of plasma membranes models

We set to explore how universal and quantitatively consistent the correlation between PM bending rigidity and molecular order (C-Laurdan GP-value) observed in GPMV originating from U2OS cells is. To this end, NIH3T3 (fibroblast mouse embryo) and MDA-MD-231 (epithelial human metastatic) cells were cultured and GPMVs extracted by incubation with DTT/PFA (see Methods and **SI** **Fig. S4**). Even though all cell models considered here were of different phenotype and morphology, their bending rigidity and C-Laurdan GP values are well fit by a single linear relation (**Fig. 3a**). Additionally, literature data for membranes composed of red blood cell lipid extracts, influenza virus particles (which bud from highly ordered plasma membrane domains) and *in-vitro* Dictyostelium cells were considered (**Fig. 3b**). Extrapolation of the linear correlation observed for the three cell lines explored here, fits the literature data for various PM models surprisingly well (**Fig. 3b**). The seemingly universal correlation calls for a rather general mechanism determining the relation between C-Laurdan GP values and bending rigidity in PM membranes.

In the commonly used elastic sheet model for bilayer bending, the bending rigidity scales as *κ*~*K*_*A*_*d*^2^ with chemical composition reflected in the material constant *K*_*A*_, which is the stretching elasticity constant, and bilayer thickness *d*, characterizing the bilayer structure. Because of the quadratic dependence, it is tempting to assign a dominant role of the membrane thickness on the bending rigidity variations. Indeed, GP values also scale with membrane thickness, which is generally also a measure of membrane lipid order. However, what remains unclear is why and under which conditions the empirically defined GP value would show approximately the same ~*d*^2^ scaling. An additional challenge is the chemical heterogeneity of lipids, which can be better understood by comparing our data to well-characterized synthetic lipid bilayers.

### Comparison to synthetic lipid membranes

Interestingly, in synthetic membranes, correlation between order, bending rigidity and cholesterol content was previously thought to be universal with cholesterol attributing stiffness and membrane thickening, see e.g. (59). However, more recent studies revealed lipid specific effects to membrane mechanics and molecular order (23, 60, 61). For example, in synthetic lipid membranes made of POPC:Chol, lipid order (as measured by GP-value) and bending rigidity are strongly positively correlated, while in DOPC:Chol and SM:Chol membranes they show no significant correlation **(Fig. 3b)**; here POPC refers to palmitoyloleoylphosphatidylcholine, DOPC to dioleoylphatidylphocholine and SM to sphingomyelin. Similarly, membrane properties in more complex lipid compositions follow intuitive “universal” trends only along certain compositional trajectories (62). Notably, bending rigidity and GP values of PM extracts studied here seem to follow a trend similar to that of POPC membranes with varying cholesterol fractions. It remains to be seen if this reflects the abundance of PC lipids with single-chain unsaturation and cholesterol in the plasma membrane (13). Note that absolute GP values and bending rigidities might vary depending on the exact measurement conditions, e.g. selection of the probe and spectral windows for GP calculation buffer conditions. For example literature GP values for pure DOPC vary form −0.35 (56), and −0.38 (57) to −0.5 (58). Such deviations will skew the diagram shown and numerical values of the κ(GP) fit in **Fig. 3b**.

In absolute value, the GPMV data for bending rigidity and GP are closest to those of bilayers composed of DOPC:Chol at high (>30%) cholesterol fraction **(Fig. 3b)**. However, it should be noted that a large fraction of GPMVs, mainly when isolated at sparse cell growth, exhibit a bending rigidity below 20*k*_*B*_*T*, which is lower than typical values of lipid membranes with a single *cis*-unsaturation in the acyl chains. Bending rigidities down to 10*k*_*B*_*T* were measured for membranes made of lipids with poly-unsaturated fatty acids (63) and indeed polyunsaturated lipids are abundant in the PM and GPMVs (13). Interestingly, also ternary mixtures of DOPC:SM:Chol fall into similar range of GPMV GP value and bending rigidity, while ternary mixtures with lower degree of lipid acyl chain unsaturation, e.g. SOPC:SM:Chol, appear to be stiffer (64). Extraction of 60% PM cholesterol softened GPMV membranes by about 30% which by absolute value is relatively little compared to the cholesterol dependence of POPC membranes, but represents a considerably stronger stiffening effect due to cholesterol than for lipids with higher degree of unsaturation as shown for DOPC:Chol **(Fig. 3b)**. (Poly)unsaturated lipids may act as a *buffer* to damp down variations in PM stiffness and in this way, contribute to the homeostasis of the cell. This might be one reason why cells invest energy into the synthesis and repair of these oxidation-prone lipids (65).

Another contribution to the low values of bending rigidities of GPMVs might be the high membrane protein content (about 40% by mass is contained in GPMVs (66)), and proteins and peptides are often observed to reduce membrane bending rigidity (20, 67–69). GPMV membrane softening by transmembrane proteins is also consistent with lower bending rigidity values (κ≈10*k*_*B*_*T*) found in GPMVs derived from HEK cells overexpressing membrane protein (15).

### Conclusion

GPMVs, which retain most of the compositional complexity of native PM, seem to be compatible with the wide range of biophysical tools developed for synthetic giant unilamellar vesicles (5, 7). This makes GPMVs an interesting model system to study PM elasticity. We found that GPMV bending deformation can be described by a single parameter, the bending rigidity, the values of which are lower than for most of those obtained on synthetic lipid vesicles. This outcome should facilitate modelling and simulation studies of PM-like membranes, and serve for comparison with synthetic lipid bilayers.

It is important to note that the inherent PM lipid asymmetry is not fully preserved in GPMVs (in particular the asymmetry resulting from flipping of phosphatidylserine lipids) and certain signalling lipids (such as PIP2) might be lost (70). Even though studies on model membranes have attempted to characterise the effect of asymmetry on membrane phase state and mechanics (71–73) the effects on bending rigidity resulting from partial loss of membrane asymmetry are difficult to predict. This is partly because bilayer bending rigidity cannot be considered as a simple superposition of the bending rigidities of the composing monolayers (74). Previous studies have indicated large stiffening effects associated with membrane phospholipid asymmetry (75, 76), but these effects could be compensated by fast-flipping membrane components such as cholesterol (77), which is abundant in GPMVs. Because we cannot yet control the extent of GPMV lipid asymmetry this questions is not directly addressed here.

Even though we have explored only three cell lines and can only speculate for universality of behaviour over any cell type, our results strongly suggest that external chemical and morphological stimuli, which have been previously reported to influence lipid order of cellular PM, also directly affect elasticity of PM-derived GPMVs, which points to a functional role of PM membrane stiffness or order. Potential roles of PM bending rigidity variation include cellular particle uptake or generation of exocytotic vesicles (2) and regulation of receptor signalling via membrane fluctuations (15, 78). In the future, it would be interesting to investigate which signaling pathways modulate these membrane properties.

One important result of our work demonstrates that in GPMVs membrane structure (GP value, order) and viscosity, probed at molecular level by C-Laurdan and fluorescent rotors, are correlated to PM mechanical properties on a larger scale. This extends the use of C-Laurdan and molecular rotors as non-invasive tools to study elastic properties of PM in live cells, decoupled from the mechanical influence of cytoskeleton. It remains to be seen, whether the correlation between GP value and bending rigidity holds also for organelle membranes which are otherwise hard to probe directly. Further studies on the relationship between the compositional complexity and membrane mechanical properties will give invaluable information on the physiology of cellular membranes.

Finally, it should be noted, that the stable correlation between membrane order, dynamic microviscosity and equilibrium mechanical parameter of bending rigidity indicates a possible pathway for cells to sense and regulate membrane compositional variations via mechanosensing (79).

## Material and Methods

### GPMV Isolation

Before experiments, U2OS, NIH 3T3 or MDA-MD-231 cells (obtained from ATTC) were plated under identical conditions in T-25 culture flaks and cultured in DMEM medium supplement with 10% FBS (Sigma F7524) and 1% Pen Strep (Thermo Fisher Scientific). For GPMV isolation at varying cell densities, cells were harvested from a confluent T-25 culture flask (about 2.7 · 10^6^ cells) and re-seeded in T-25 flasks after 1:10, 1:5 and 1:2 dilution in medium. After incubation for 24 hours, GPMVs were isolated according to the protocol reported in (22). U2OS cells were incubated with 2 mM DTT, 25 mM PFA at 37°C for 1 hour. Sometimes cells were labeled before GPMV extraction, using the membrane dye Fast-DiIC18 (Thermo Fisher Scientific) by incubation of cells together with dye in PBS buffer for 10 minutes at 4°C. Experiments were generally conducted in triplicates except those at varying growth density and cholesterol extraction, which were conducted in duplicates.

### Fluctuation spectroscopy

GPMVs were extracted in nearly isoosmolar conditions (320 mOsmol/L, see buffer composition above). To allow for optically resolvable membrane fluctuations, the osmolality of the solution surrounding the GPMVs was slowly increased by evaporation. A 30 μl drop of GPMV suspension was left on a cover glass for 5 minutes before a chamber was formed by a top cover glass and “press-to-seal” silicon isolators (Sigma-Aldrich). Cover glasses were coated with BSA solution to suppress unspecific adhesion: cover glasses cleaned with ethanol were incubated for 30 minutes in 10 mg/ml BSA solution (fatty acid free, A1595 Sigma-Aldrich) and rinsed with double distilled water before usage.

The majority of defect-free GPMVs underwent optically resolvable fluctuations. Membrane bending rigidity was then measured by fluctuation analysis of the thermally induced motion of the membrane. Details of the method are published elsewhere (23). Experiments were performed on an Axio Observer D1 microscope (Zeiss, Germany) using a 40× objective in phase contrast mode. Imaging was done using a low noise liquid-cooled digital camera pco.edge 5.5. We acquired a total of 2000-4000 snapshots per vesicle with exposure time of 200 μs. Frame rates were varied between 20 and 100 frames per second with no difference in measured bending rigidity within the statistical uncertainty.

For a quasi-spherical vesicle of radius *R*_*GPMG*_, the dimensionless mean square amplitudes of the spherical harmonic modes behave as

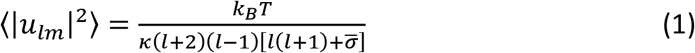

where 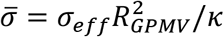 is the rescaled effective tension, which together with the bending rigidity κ s treated as a fit parameter.

To check for artefacts in the fluctuation spectra resulting from gravity-induced vesicle deformation, the criterion developed in (80) was employed. The density of the GPMV cytosol-like interior was estimated by assuming a macromolecular volume fraction of 20 wt% with a density of 1.3 g/ml (81) and a density of 1 g/ml of the outside buffer. The criterion by Henriksen et al. (80) predicts negligible deformation for the typically small GPMVs with radii below 10 μm and tension values estimated from the fit using Eq. (1).

### Modulation of cholesterol levels in cells

Cell-culture grade methyl-β-cyclodextrin (MβCD, Sigma) was either used as obtained from the manufacturer or pre-complexed with saturating concertation of cholesterol in sterile conditions (82). Cells were washed and incubated with 10 mM of empty or cholesterol-loaded MβCD in DMEM buffer for 25 min at 37°C. Control was incubated with DMEM buffer only.

### Measurements at elevated temperature

GPMV were extracted as described above using DTT/PFA chemicals at 70% confluency. Sample temperature was controlled using a home build chamber described in detail elsewhere (83). Fluctuation analysis was performed as described above.

### GP and FLIM imaging

GPMV samples were split in half and labelled at room temperature with either C-Laurdan (0.4 μM) or molecular rotor (compound 1 in (30), 0.05 μg/ml) for 30 minutes (22). Although it is a relatively new probe, C-Laurdan is reported to respond to solvent polarity similarly to Laurdan (84–86). Molecular rotors, on the other hand, change their lifetime depending on the viscosity of the environment (30, 87). For GPMVs labelled with C-Laurdan, spectral imaging was carried out with confocal microscope Zeiss 780 using 40× water immersion objective (NA 1.2) (27). C-Laurdan was excited at 405 nm and the emission was collected with a multichannel spectral detector in the range 410–540 nm, with a spectral step of 8.9 nm per channel. GP maps were calculated with the Fiji plugin freely available (27).

For FLIM imaging of the molecular rotor, labelled vesicles were transferred into glass-bottom 8-well microscopy slides (ibidi) and placed on an inverted laser-scanning confocal microscope (Leica SP8 STED) equipped with a time-correlated single photon counting (TCSPC) module (PicoQuant) and a 63x water immersion objective. Fluorescence in the equatorial plane of GPMVs was excited by a white light laser tuned to 488 nm and emission collected by a hybrid detector in the range 500–560 nm at a 160-nm pixel size. The photon streams were recorded until 300 photons per channel were detected. Average lifetime values for each GPMV were then extracted by fitting a bi-exponential decay curve to the histogram of data collected from pixels within the masked area along the perimeter of the GPMV, using PicoQuant SymphoTime64 software. As noted previously (87), the fast decay component of this probe in membranes (in our case with lifetimes around 0.6–0.8 ns and relative intensities around 20–30%) originates from a fraction of molecules that are insensitive to membrane viscosity due to their conformation (e.g. lying in the membrane plane). The values of fit component with longer lifetime were therefore used for further comparison.

## Acknowledgments

JS would like to acknowledge Dennis Discher for providing the USO2 cell line, Christine Pilz-Allen and Amaia Cipitria for kind donation of NIH-3T3 and MDA-MD-231 cells, and Eleanor Ewins for critical reading of the manuscript, ES acknowledges Prof. Marina Kuimova (Imperial Collage London) for the molecular rotor viscosity probe. We would like to thank Wolfson Imaging Centre for providing imaging tools. The authors also thank their funding: I.U. is financed by Marie Skłodowska-Curie fellowship (grant no. 707348). E.S. is supported by EMBO long term (ALTF 636-2013), Marie Skłodowska-Curie Intra-European Fellowships (MEMBRANE DYNAMICS-627088) and Newton-Katip Celebi Institutional Links grant (352333122). This work is supported by the Wolfson Foundation (ref. 18272), the Medical Research Council (MRC, grant number MC_UU_12010/unit programmes G0902418 and MC_UU_12025), MRC/BBSRC/ESPRC (grant number MR/K01577X/1), the Wellcome Trust (grant ref. 104924/14/Z/14), Deutsche Forschungsgemeinschaft (Research unit 1905 “Structure and function of the peroxisomal translocon”), and internal University of Oxford funding (EPA Cephalosporin Fund and John Fell Fund). This work is part of the MaxSynBio consortium which was jointly funded by the Federal Ministry of Education and Research of Germany and the Max Planck Society.

## Author contributions

JS, ES and RD designed the experiments, JS ES and IU conducted the experiments and analysed the data, and all authors wrote the manuscript.

## Supporting Information

**Fig. S1.**
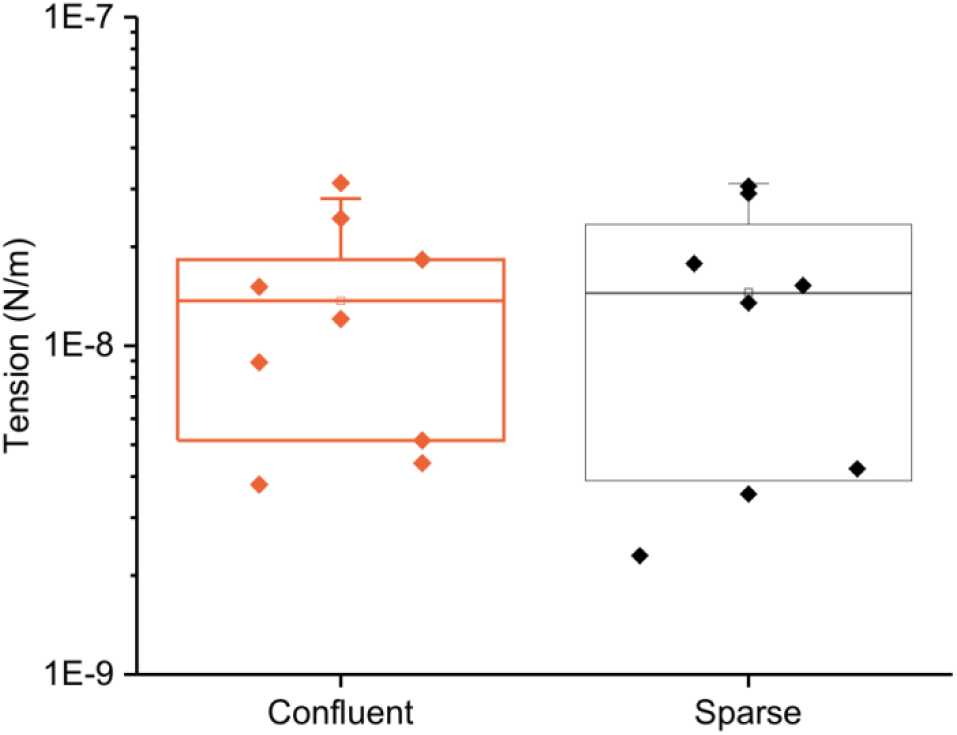
Typical membrane tension values measured by fluctuation spectroscopy. The two samples correspond to a subset of GPMVs from Fig. 2b and indicate the cell confluency at GPMV isolation (GPMVs were extracted by DTT/PFA). Each data point indicates one individual vesicle. Boxes have the conventional meaning of lower 25% and 75% quartile around the population mean value (middle line) and error bars indicate 1.5 std. dev.

**Fig. S2.**
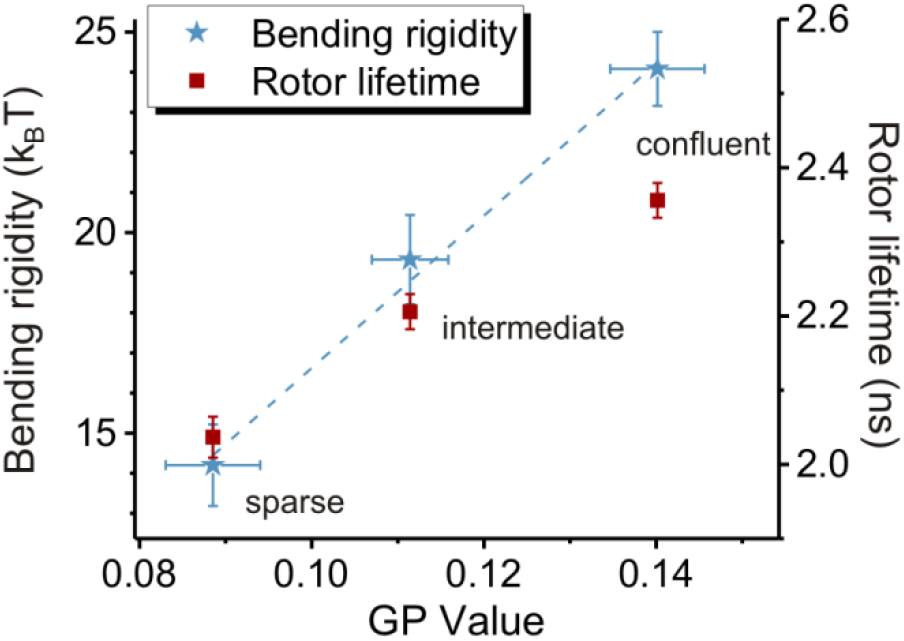
Correlation between values of bending rigidity, GP value and viscosity measured on U2OS cells at varying cell density. Error bars indicate std. error from mean (n=9,17,20 GPMVs).

**Fig. S3.**
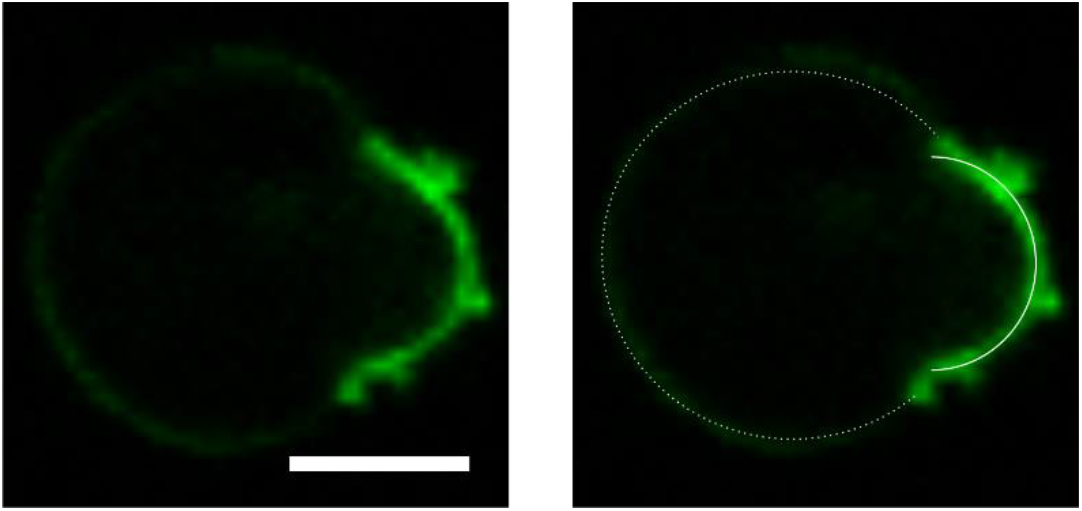
Confocal cross section obtained on a phase separated GPMV. At room temperature the majority of GPMVs exhibits one single fluid phase. However, a fraction of GPMVs was found to be phase-separated into two liquid phases. Fast-DiIC18 (dye) partitions into the liquid disordered phase (shown in green). In the right image the curvature of the two membrane segments is shown with solid and dotted contours. As it can be seen from their intersection, the liquid disordered phase (domain on the right side) deforms to match the curvature of the liquid ordered phase, indicating that it is energetically more favourable to bend the liquid discorded domain. Scale bar indicates 5μm.

**Fig. S4.**
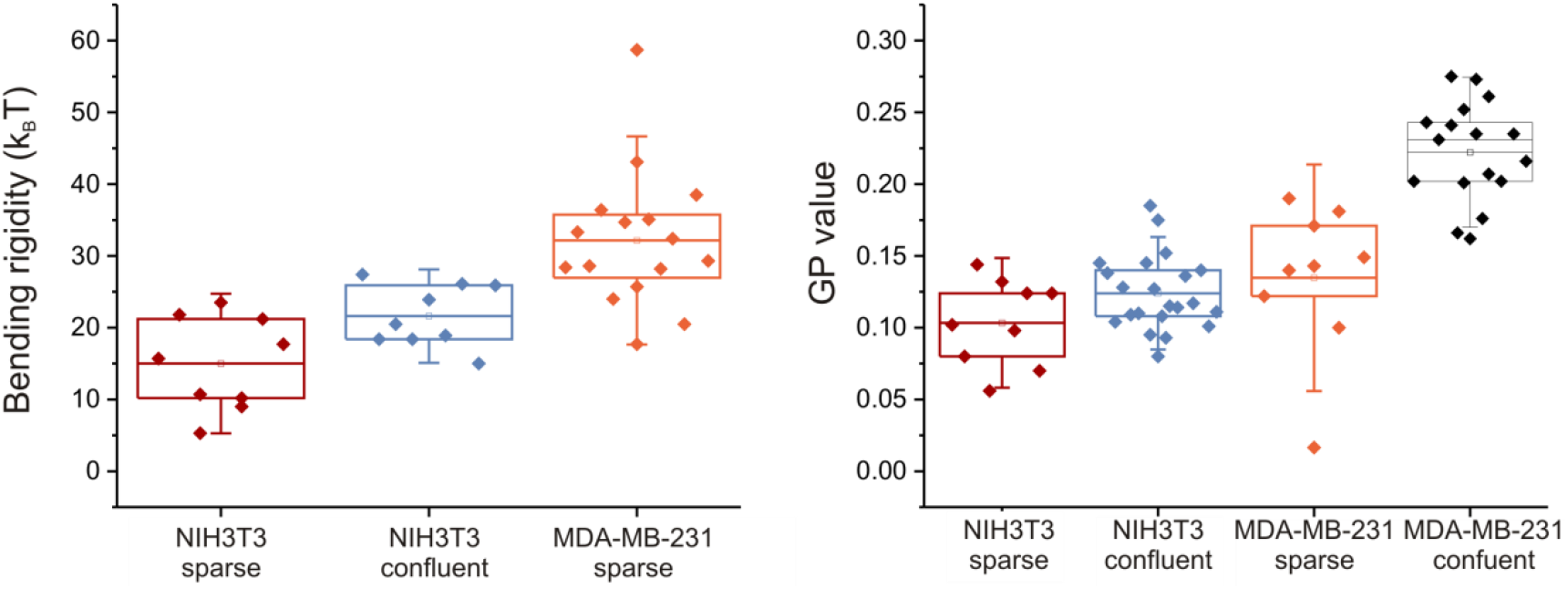
Effect of cell confluency on the mechanical and molecular properties of three cell lines. Each data point represents one measurement on a single GPMV. GPMVs were isolated using 2 mM DTT + 25 mM PFA. Boxes have the conventional meaning of lower 25% and 75% quartile around the population mean value (middle line) and error bars indicate 1.5 std. dev. Bending rigidity data on MDA-MD-231 cells at spare conditions could not be obtained due to low yield of suitable GPMVs.

## References

1. Jarsch IK, Daste F, Gallop JL. Membrane curvature in cell biology: An integration of molecular mechanisms. The Journal of Cell Biology. 2016;214(4):375–87.

2. Agudo-Canalejo J, Lipowsky R. Critical Particle Sizes for the Engulfment of Nanoparticles by Membranes and Vesicles with Bilayer Asymmetry. Acs Nano. 2015;9(4):3704–20.

3. Stachowiak JC, Brodsky FM, Miller EA. A cost–benefit analysis of the physical mechanisms of membrane curvature. Nat Cell Biol. 2013;15(9):1019.

4. Muro E, Atilla-Gokcumen GE, Eggert US. Lipids in cell biology: how can we understand them better? Mol Biol Cell. 2014;25(12):1819–23.

5. Dimova R, Said A, Natalya B, Vesselin N, Karin AR, Reinhard L. A practical guide to giant vesicles. Probing the membrane nanoregime via optical microscopy. Journal of Physics: Condensed Matter. 2006;18(28):S1151.

6. Walde P, Cosentino K, Engel H, Stano P. Giant Vesicles: Preparations and Applications. ChemBioChem. 2010;11(7):848–65.

7. Dimova R. Giant Vesicles and Their Use in Assays for Assessing Membrane Phase State, Curvature, Mechanics, and Electrical Properties. Annual review of biophysics. 2019;48.

8. Scott RE. Plasma membrane vesiculation: a new technique for isolation of plasma membranes. Science. 1976;194(4266):743.

9. Baumgart T, Hammond AT, Sengupta P, Hess ST, Holowka DA, Baird BA, et al. Large-scale fluid/fluid phase separation of proteins and lipids in giant plasma membrane vesicles. Proceedings of the National Academy of Sciences. 2007;104(9):3165–70.

10. Veatch SL, Cicuta P, Sengupta P, Honerkamp-Smith A, Holowka D, Baird B. Critical fluctuations in plasma membrane vesicles. ACS chemical biology. 2008;3(5):287–93.

11. Braig S, Schmidt BS, Stoiber K, Händel C, Möhn T, Werz O, et al. Pharmacological targeting of membrane rigidity: implications on cancer cell migration and invasion. New J Phys. 2015;17(8):083007.

12. Bauer B, Davidson M, Orwar O. Proteomic Analysis of Plasma Membrane Vesicles. Angew Chem Int Ed. 2009;48(9):1656–9.

13. Levental I, Veatch SL. The Continuing Mystery of Lipid Rafts. J Mol Biol. 2016;428(24, Part A):4749–64.

14. Özhan G, Sezgin E, Wehner D, Pfister AS, Kühl SJ, Kagermeier-Schenk B, et al. Lypd6 enhances Wnt/β-catenin signaling by promoting Lrp6 phosphorylation in raft plasma membrane domains. Developmental cell. 2013;26(4):331–45.

15. Steinkühler J, Różycki B, Alvey C, Lipowsky R, Weikl TR, Dimova R, et al. Membrane fluctuations and acidosis regulate cooperative binding of “marker of self” CD47 with macrophage checkpoint receptor SIRPα. Journal of Cell Science. 2018.

16. Schneider F, Waithe D, Clausen MP, Galiani S, Koller T, Ozhan G, et al. Diffusion of lipids and GPI-anchored proteins in actin-free plasma membrane vesicles measured by STED-FCS. Mol Biol Cell. 2017;28(11):1507–18.

17. Dai J, Sheetz MP. Membrane Tether Formation from Blebbing Cells. Biophys J. 1999;77(6):3363–70.

18. Byfield FJ, Aranda-Espinoza H, Romanenko VG, Rothblat GH, Levitan I. Cholesterol Depletion Increases Membrane Stiffness of Aortic Endothelial Cells. Biophys J. 2004;87(5):3336–43.

19. Pontes B, Ayala Y, Fonseca ACC, Romao LF, Amaral RF, Salgado LT, et al. Membrane Elastic Properties and Cell Function. Plos One. 2013;8(7).

20. Dimova R. Recent developments in the field of bending rigidity measurements on membranes. Advances in Colloid and Interface Science. 2014;208:225–34.

21. Charras GT, Coughlin M, Mitchison TJ, Mahadevan L. Life and Times of a Cellular Bleb. Biophys J. 2008;94(5):1836–53.

22. Sezgin E, Kaiser H-J, Baumgart T, Schwille P, Simons K, Levental I. Elucidating membrane structure and protein behavior using giant plasma membrane vesicles. Nat Protocols. 2012;7(6):1042–51.

23. Gracia RS, Bezlyepkina N, Knorr RL, Lipowsky R, Dimova R. Effect of cholesterol on the rigidity of saturated and unsaturated membranes: fluctuation and electrodeformation analysis of giant vesicles. Soft Matter. 2010;6(7):1472–82.

24. Shi Z, Graber ZT, Baumgart T, Stone HA, Cohen AE. Cell Membranes Resist Flow. Cell. 2018;175(7):1769–79.e13.

25. Colom A, Derivery E, Soleimanpour S, Tomba C, Molin MD, Sakai N, et al. A fluorescent membrane tension probe. Nature Chemistry. 2018;10(11):1118–25.

26. Levental I, Grzybek M, Simons K. Raft domains of variable properties and compositions in plasma membrane vesicles. Proceedings of the National Academy of Sciences. 2011;108(28):11411–6.

27. Sezgin E, Waithe D, Bernardino de la Serna J, Eggeling C. Spectral imaging to measure heterogeneity in membrane lipid packing. ChemPhysChem. 2015;16(7):1387–94.

28. Parasassi T, Krasnowska EK, Bagatolli L, Gratton E. Laurdan and Prodan as Polarity-Sensitive Fluorescent Membrane Probes. Journal of Fluorescence. 1998;8(4):365–73.

29. Sezgin E, Sadowski T, Simons K. Measuring Lipid Packing of Model and Cellular Membranes with Environment Sensitive Probes. Langmuir. 2014;30(27):8160–6.

30. Lopez-Duarte I, Vu TT, Izquierdo MA, Bull JA, Kuimova MK. A molecular rotor for measuring viscosity in plasma membranes of live cells. Chemical Communications. 2014;50(40):5282–4.

31. Kubánková M, López-Duarte I, Kiryushko D, Kuimova MKJSm. Molecular rotors report on changes in live cell plasma membrane microviscosity upon interaction with beta-amyloid aggregates. 2018;14(46):9466–74.

32. Gerstle R, Desai R, Veatch S. Dithiothreitol Raises Transition Temperatures in Giant Plasma Membrane Vesicles. Biophys J. 2017;112(3):519a.

33. Levental I, Lingwood D, Grzybek M, Coskun Ü, Simons K. Palmitoylation regulates raft affinity for the majority of integral raft proteins. Proceedings of the National Academy of Sciences. 2010;107(51):22050–4.

34. Amaro M, Reina F, Hof M, Eggeling C, Sezgin E. Laurdan and Di-4-ANEPPDHQ probe different properties of the membrane. Journal of Physics D: Applied Physics. 2017;50(13):134004.

35. Snijder B, Sacher R, Rämö P, Damm E-M, Liberali P, Pelkmans L. Population context determines cell-to-cell variability in endocytosis and virus infection. Nature. 2009;461(7263):520–3.

36. Frechin M, Stoeger T, Daetwyler S, Gehin C, Battich N, Damm E-M, et al. Cell-intrinsic adaptation of lipid composition to local crowding drives social behaviour. Nature. 2015;523(7558):88–91.

37. Noutsi P, Gratton E, Chaieb SJPo. Assessment of membrane fluidity fluctuations during cellular development reveals time and cell type specificity. 2016;11(6):e0158313.

38. Gray EM, Díaz-Vázquez G, Veatch SL. Growth conditions and cell cycle phase modulate phase transition temperatures in RBL-2H3 derived plasma membrane vesicles. Plos One. 2015;10(9):e0137741.

39. Gaus K, Gratton E, Kable EPW, Jones AS, Gelissen I, Kritharides L, et al. Visualizing lipid structure and raft domains in living cells with two-photon microscopy. Proc Natl Acad Sci U S A. 2003;100(26):15554–9.

40. Levental I, Byfield FJ, Chowdhury P, Gai F, Baumgart T, Janmey PA. Cholesterol-dependent phase separation in cell-derived giant plasma-membrane vesicles. The Biochemical journal. 2009;424(2):163–7.

41. Kilsdonk EPC, Yancey PG, Stoudt GW, Bangerter FW, Johnson WJ, Phillips MC, et al. Cellular Cholesterol Efflux Mediated by Cyclodextrins. J Biol Chem. 1995;270(29):17250–6.

42. Yancey PG, Rodrigueza WV, Kilsdonk EPC, Stoudt GW, Johnson WJ, Phillips MC, et al. Cellular Cholesterol Efflux Mediated by Cyclodextrins: DEMONSTRATION OF KINETIC POOLS AND MECHANISM OF EFFLUX. J Biol Chem. 1996;271(27):16026–34.

43. Zidovetzki R, Levitan I. Use of cyclodextrins to manipulate plasma membrane cholesterol content: evidence, misconceptions and control strategies. Biochimica et biophysica acta. 2007;1768(6):1311–24.

44. Baumgart T, Das S, Webb WW, Jenkins JT. Membrane elasticity in giant vesicles with fluid phase coexistence. Biophys J. 2005;89(2):1067–80.

45. Gutlederer E, Gruhn T, Lipowsky R. Polymorphism of vesicles with multi-domain patterns. Soft Matter. 2009;5(17):3303–11.

46. Celli A, Gratton E. Dynamics of lipid domain formation: Fluctuation analysis. Biochimica et Biophysica Acta (BBA)-Biomembranes. 2010;1798(7):1368–76.

47. Carravilla P, Nieva JL, Gonñi FlM, Requejo-Isidro J, Huarte N. Two-photon Laurdan studies of the ternary lipid mixture DOPC: SM: cholesterol reveal a single liquid phase at sphingomyelin: cholesterol ratios lower than 1. Langmuir. 2015;31(9):2808–17.

48. Kulig W, Jurkiewicz P, Olżyńska A, Tynkkynen J, Javanainen M, Manna M, et al. Experimental determination and computational interpretation of biophysical properties of lipid bilayers enriched by cholesteryl hemisuccinate. Biochimica et Biophysica Acta (BBA)-Biomembranes. 2015;1848(2):422–32.

49. Kaiser H-J, Surma MA, Mayer F, Levental I, Grzybek M, Klemm RW, et al. Molecular Convergence of Bacterial and Eukaryotic Surface Order. J Biol Chem. 2011;286(47):40631–7.

50. Sezgin E, Gutmann T, Buhl T, Dirkx R, Grzybek M, Coskun Ü, et al. Adaptive lipid packing and bioactivity in membrane domains. Plos One. 2015;10(4):e0123930.

51. Henriksen J, Rowat AC, Ipsen JH. Vesicle fluctuation analysis of the effects of sterols on membrane bending rigidity. Eur Biophys J. 2004;33(8):732–41.

52. Mazeres S, Joly E, Lopez A, Tardin C. Characterization of M-laurdan, a versatile probe to explore order in lipid membranes. F1000Research. 2014;3.

53. Gerl MJ, Sampaio JL, Urban S, Kalvodova L, Verbavatz J-M, Binnington B, et al. Quantitative analysis of the lipidomes of the influenza virus envelope and MDCK cell apical membrane. J Cell Biol. 2012;196(2):213–21.

54. Schaap IA, Eghiaian F, des Georges A, Veigel C. Effect of envelope proteins on the mechanical properties of influenza virus. J Biol Chem. 2012;287(49):41078–88.

55. Simson R, Wallraff E, Faix J, Niewöhner J, Gerisch G, Sackmann EJBj. Membrane bending modulus and adhesion energy of wild-type and mutant cells of Dictyostelium lacking talin or cortexillins. 1998;74(1):514–22.

56. Kim HM, Kim BR, Choo HJ, Ko YG, Jeon SJ, Kim CH, et al. Two-Photon Fluorescent Probes for Biomembrane Imaging: Effect of Chain Length. ChemBioChem. 2008;9(17):2830–8.

57. Kaiser H-J, Lingwood D, Levental I, Sampaio JL, Kalvodova L, Rajendran L, et al. Order of lipid phases in model and plasma membranes. Proceedings of the National Academy of Sciences. 2009;106(39):16645–50.

58. Sáenz JP, Sezgin E, Schwille P, Simons K. Functional convergence of hopanoids and sterols in membrane ordering. Proceedings of the National Academy of Sciences. 2012;109(35):14236–40.

59. Henriksen J, Rowat AC, Brief E, Hsueh Y, Thewalt J, Zuckermann M, et al. Universal behavior of membranes with sterols. Biophys J. 2006;90(5):1639–49.

60. Pan J, Mills TT, Tristram-Nagle S, Nagle JF. Cholesterol perturbs lipid bilayers nonuniversally. Phys Rev Lett. 2008;100(19):198103.

61. M’Baye G, Mély Y, Duportail G, Klymchenko ASJBj. Liquid ordered and gel phases of lipid bilayers: fluorescent probes reveal close fluidity but different hydration. 2008;95(3):1217–25.

62. Bleecker Joan V, Cox Phillip A, Keller Sarah L. Mixing Temperatures of Bilayers Not Simply Related to Thickness Differences between Lo and Ld Phases. Biophys J. 2016;110(11):2305–8.

63. Rawicz W, Olbrich KC, McIntosh T, Needham D, Evans E. Effect of chain length and unsaturation on elasticity of lipid bilayers. Biophys J. 2000;79(1):328–39.

64. Rawicz W, Smith B, McIntosh T, Simon S, Evans E. Elasticity, strength, and water permeability of bilayers that contain raft microdomain-forming lipids. Biophys J. 2008;94(12):4725–36.

65. Manni MM, Tiberti ML, Pagnotta S, Barelli H, Gautier R, Antonny B. Acyl chain asymmetry and polyunsaturation of brain phospholipids facilitate membrane vesiculation without leakage. eLife. 2018;7:e34394.

66. Seeliger J, Erwin N, Rosin C, Kahse M, Weise K, Winter R. Exploring the structure and phase behavior of plasma membrane vesicles under extreme environmental conditions. Physical Chemistry Chemical Physics. 2015;17(11):7507–13.

67. Fowler PW, Hélie J, Duncan A, Chavent M, Koldsø H, Sansom MS. Membrane stiffness is modified by integral membrane proteins. Soft Matter. 2016;12(37):7792–803.

68. Shchelokovskyy P, Tristram-Nagle S, Dimova R. Effect of the HIV−1 fusion peptide on the mechanical properties and leaflet coupling of lipid bilayers. New J Phys. 2011;13(2):025004.

69. Sorkin R, Huisjes R, Bošković F, Vorselen D, Pignatelli S, Ofir-Birin Y, et al. Nanomechanics of Extracellular Vesicles Reveals Vesiculation Pathways. Small. 2018;14(39):1801650.

70. Keller H, Lorizate M, Schwille P. PI (4, 5) P2 Degradation Promotes the Formation of Cytoskeleton-Free Model Membrane Systems. Chemphyschem. 2009;10(16):2805–12.

71. Kubsch B, Robinson T, Lipowsky R, Dimova R. Solution Asymmetry and Salt Expand Fluid-Fluid Coexistence Regions of Charged Membranes. Biophys J. 2016;110(12):2581–4.

72. Dasgupta R, Miettinen MS, Fricke N, Lipowsky R, Dimova R. The glycolipid GM1 reshapes asymmetric biomembranes and giant vesicles by curvature generation. Proceedings of the National Academy of Sciences. 2018;115(22):5756–61.

73. Karimi M, Steinkühler J, Roy D, Dasgupta R, Lipowsky R, Dimova R. Asymmetric ionic conditions generate large membrane curvatures. Nano letters. 2018;18(12):7816–21.

74. Sreekumari A, Lipowsky RJTJocp. Lipids with bulky head groups generate large membrane curvatures by small compositional asymmetries. 2018;149(8):084901.

75. Lu L, Doak WJ, Schertzer JW, Chiarot PR. Membrane mechanical properties of synthetic asymmetric phospholipid vesicles. Soft Matter. 2016;12(36):7521–8.

76. Elani Y, Purushothaman S, Booth PJ, Seddon JM, Brooks NJ, Law RV, et al. Measurements of the effect of membrane asymmetry on the mechanical properties of lipid bilayers. Chemical Communications. 2015;51(32):6976–9.

77. Miettinen MS, Lipowsky R. Bilayer membranes with frequent flip-flops have tensionless leaflets. Nano Letters. 2019.

78. Krobath H, Różycki B, Lipowsky R, Weikl TR. Binding cooperativity of membrane adhesion receptors. Soft Matter. 2009;5(17):3354–61.

79. Dymond MK. Mammalian phospholipid homeostasis: homeoviscous adaptation deconstructed by lipidomic data driven modelling. Chem Phys Lipids. 2015;191:136–46.

80. Henriksen JR, Ipsen JH. Thermal undulations of quasi-spherical vesicles stabilized by gravity. Eur Phys J E. 2002;9(4):365–74.

81. Görisch SM, Lichter P, Rippe K. Mobility of multi-subunit complexes in the nucleus: accessibility and dynamics of chromatin subcompartments. Histochemistry and cell biology. 2005;123(3):217–28.

82. Levitan I, Christian AE, Tulenko TN, Rothblat GH. Membrane cholesterol content modulates activation of volume-regulated anion current in bovine endothelial cells. The Journal of general physiology. 2000;115(4):405–16.

83. Kubsch B, Robinson T, Steink hler J, Dimova R. Phase Behavior of Charged Vesicles Under Symmetric and Asymmetric Solution Conditions Monitored with Fluorescence Microscopy. 2017(128):e56034.

84. Kim HM, Choo HJ, Jung SY, Ko YG, Park WH, Jeon SJ, et al. A two-photon fluorescent probe for lipid raft imaging: C-laurdan. ChemBioChem. 2007;8(5):553–9.

85. Barucha-Kraszewska J, Kraszewski S, Ramseyer C. Will C-Laurdan dethrone Laurdan in fluorescent solvent relaxation techniques for lipid membrane studies? Langmuir. 2013;29(4):1174–82.

86. Mazeres S, Fereidouni F, Joly E. Using spectral decomposition of the signals from laurdan-derived probes to evaluate the physical state of membranes in live cells. F1000Research. 2017;6.

87. Wu Y, Štefl M, Olzyńska A, Hof M, Yahioglu G, Yip P, et al. Molecular rheometry: direct determination of viscosity in L o and L d lipid phases via fluorescence lifetime imaging. Physical Chemistry Chemical Physics. 2013;15(36):14986–93.

